# The effects of arbuscular mycorrhizal fungi (AMF) and *Rhizophagus irregularis* on soil microorganisms assessed by metatranscriptomics and metaproteomics

**DOI:** 10.1101/860932

**Authors:** D.R. Lammel, D. Meierhofer, P. Johnston, S. Mbedi, M.C. Rillig

## Abstract

Arbuscular mycorrhizal fungi (AMF) form symbioses with approximately 80% of plant species and potentially benefit their hosts (e.g. nutrient acquisition) and the soil environment (e.g. soil aggregation). AMF also affect soil microbiota and soil multifunctionality. We manipulated AMF presence (via inoculation of non-sterile soil with *Rhizophagus irregularis* and using a hyphal compartment design) and used RNA-seq and metaproteomics to assess AMF roles in soil. The results indicated that AMF drove an active soil microbial community expressing transcripts and proteins related to nine metabolic functions, including the metabolism of C and N. We suggest two possible mechanisms: 1) the AMF hyphae produce exudates that select a beneficial community, or, 2) the hyphae compete with other soil microbes for available nutrients and consequently induce the community to mineralize nutrients from soil organic matter. We also identified candidate proteins that are potentially related to soil aggregation, such as Lpt and HSP60. Our results bridge microbial ecology and ecosystem functioning. We show that the AMF hyphosphere contains an active community related to soil respiration and nutrient cycling, thus potentially improving nutrient mineralization from soil organic matter and nutrient supply to the plants.

## 1. Introduction

Recent studies indicated that plant roots select beneficial microbial communities related to nutrient cycling in the rhizosphere (Mendes et al., 2014, 2018b; Baltrus, 2017; Yan et al., 2017; Schmidt et al., 2019). Therefore, we hypothesized that arbuscular mycorrhizal fungi (AMF) also select beneficial microbial communities in their hyphosphere (the volume of soil influenced by AMF hyphae).

Previous studies have found that AMF affect the soil microbial community structure at the taxonomic level (Turrini et al., 2018; Xu et al., 2018; Bhatt et al., 2019). However, even though such diversity studies based on ribosomal DNA regions (e.g. 16/18S rRNA or ITS) are robust methods to assess microbial community structure, they usually fail to correlate to environmental processes (Hall et al., 2018; Lammel et al., 2018; Zhang et al., 2019). More accurate information related to ecosystem functioning, interactions and nutrient cycling can be obtained by assessing directly the metabolically active microbial community by RNA-seq and proteomics (Mendes et al., 2018a; Yao et al., 2018). Thus, learning about the metabolically active microbial community can be particularly important for a better understanding of arbuscular mycorrhizal fungi (AMF) interactions with the soil microbiota.

AMF are soil fungi that form a symbiosis with approximately 80% of plant species. During the symbiosis, plant roots supply these fungi with carbohydrates and fatty acids, while the fungi induce metabolic changes in the plant (including improved plant growth and better plant response to biotic and abiotic stress) (Kameoka et al.; Delavaux et al., 2017; Powell and Rillig, 2018; Begum et al., 2019; Mateus et al., 2019). In addition to intimately associating with the plant root, the fungi also grow in the soil, foraging for mineral nutrients and water, and affecting soil properties (Kameoka et al.; Rillig and Mummey, 2006; Pepe et al., 2017). Consequently, AMF have three main roles in soil, they: 1) taking up mineral nutrients and water, therefore changing soil nutrient and water dynamics; 2) interacting with and changing soil microbial communities, and 3) interacting with and modifying soil structure, e.g. by aggregating soil (Rillig and Mummey, 2006; Cavagnaro et al., 2015; Powell and Rillig, 2018; Bhatt et al., 2019).

Thus, it is not only relevant to understand how AMF change the dynamics of nutrients in the soil but also how AMF induce changes in the soil microbial community itself (Turrini et al., 2018; Bhatt et al., 2019). The inoculation of AMF, remarkably *R. irregularis*, affects soil microbial community structure at the taxonomic level (Changey et al., 2019). Moreover, some bacterial taxa are likely associated with AMF spores, for example, *Proteobacteria* and *Firmicutes* (Battini et al., 2016; Changey et al., 2019). However, detailed information about the effects of AMF on the functionally active microbial community is still unavailable.

AMF also affect soil structure and their presence and abundance in soil is typically correlated with soil aggregation (Rillig and Mummey, 2006; Schlüter et al., 2019, 2019). On the other hand, soil aggregation affects microbial communities in soil (Rillig et al., 2017; Upton et al., 2019). Soil aggregation is likely influenced by direct effects of the AMF, such as hyphal enmeshment of soil particles and production of exo-polymeric substances (EPS), and indirect effects, such as AMF effects on microbial communities and the soil food web that could then change the soil structure and aggregates (Rillig and Mummey, 2006). EPS can also be produced by the microbial community, and include mainly polysaccharides and lipopolysaccharides, which are synthesized by proteins such as Wza, Wca, Kps, Alg, and Sac, and have been reported to be potentially correlated to soil aggregation (Cania et al., 2019).

Another possible mechanism for soil aggregation could be that specific AMF proteins may act as binding agents for soil particles. In the 1990’s, a monoclonal antibody was produced (MAb32B11) for the purpose of quantifying *R. irregularis* in soil and the protein attached to that antibody was called “glomalin” (name derived from *Glomus*) and it correlated with soil aggregation (Wright et al., 1996). Later, general “glomalin related soil protein” (“GRSP”) extractions methods were proposed, but such methods were not specific and very likely biased by substances other than glomalin (Rosier et al., 2006; Gillespie et al., 2011). Thus, the initial hypotheses linking specific AMF proteins and soil aggregation are not confirmed. Gadkar and Rillig (2006) provided initial evidence that the protein that attaches to the *R. irregularis* 90’s antibody (that was first called glomalin) may be a HSP60 protein. HSP60 proteins (GroEL) are mainly molecular chaperons, but they can also have moonlight functions, and are widespread proteins present in all living organisms (Henderson et al., 2013). It is still unclear whether this protein can directly interact with soil particles, but a priority research question is to quantify the abundance of HSP60 from *R. irregularis* in soil compared to the natural occurrence of HSP60 from the overall soil microbial community.

The objective of this study was to use RNA-seq and proteomics to investigate the hypothesis that the AMF hyphosphere selects for a beneficial microbial community. We mainly investigated the AMF effects on: 1) the C, N and P soil community metabolism at the gene/protein level; 2) the soil microbial community structure itself (assessed by the taxonomic affiliation of the genes/proteins); and 3) transcripts and proteins potentially related to soil aggregation (such as the above-mentioned proteins Wza, Wca, Kps, Alg, Sac and HSP60).

## 2. Material and Methods

### Compartmentalized experimental setup with hyphal compartment

To test the effects of AMF on soil, we used the hyphal compartment approach (Johnson et al., 2001; Leifheit et al., 2014). We established two main treatments, non-sterile soil control and the same soil inoculated with *R. irregularis* (DAOM 197198) with ten replicates (ten pots) for each treatment. In all pots we placed the hyphal compartment cores, and then the control soils were further divided into two treatments: static cores and cores that were rotated three times per week. The rotation of the cores severs fungal hyphae thus avoiding the growth of AMF inside the rotated cores (Leifheit et al., 2014). So the experiment was finally divided into three operational treatments: control with static cores (C), control with rotated cores (rC) and static cores in soil inoculated with the AMF *R. irregularis* (A). Due to technical limitations of advanced molecular techniques (e.g. sequencing costs), we randomly chose four samples of each treatment for RNA-seq and five samples of each treatment for proteomics. We report all the non-molecular analysis results for all the treatments and replicates in Supplementary Material 1.

The soil used in the experiment was collected at the agricultural experimental station of the Humboldt University in Dahlem (Berlin, Germany). The top soil from a meadow (0-30 cm) was taken and air-dried at room temperature. The soil was an Albic Luvisol (World Reference Base for soil resources, 1998) and had the following physical-chemical characteristics: 73.6% sand, 18.8% silt, 7.6% clay; pH 7.1 (CaCl_2_) (analyses conducted by LUFA Rostock Agricultural Analysis and Research Institute, Germany); 1.87% C (total) and a C/N ratio 15.6 (analyzed on an Euro EA C/N analyzer, HEKAtech GmbH, Wegberg, Germany) (Rillig et al., 2010). The soil was sieved to 4mm and root and plant material removed. Twenty plastic pots (20cm diameter × 20cm high) were filled with five liters of this soil and core compartments inserted in each pot (details about the cores in Leifheit et al., 2014).

Seeds of clover (*Trifolium repens* L) were surface sterilized with 70% ethanol for 30 secs, followed by 3% sodium hypochlorite for 5 min, and then rinsed five times with autoclaved water. Ten of these seeds were sown evenly in each pot, but after germination only five plants were kept per pot. Half of the pots (ten) received the main treatment and were inoculated with *R. irregularis* and the other half, ten pots, were kept as control without inoculation. The inoculum was prepared from *R. irregularis* that was grown under *in vitro* culture conditions (Rillig et al., 2010). A piece of solid media containing hyphae and spores (“a loopful”) from the hyphae compartment (containing 20 ± 7 spores) was inoculated nearby each seed in the inoculated treatments and a “loopful” of sterile media was added nearby each seed for the controls. The plants were grown in a greenhouse (22°C/ 18°C, day/ night) and a photoperiod of 14h (a mix of natural light and supplementation of artificial light when needed). An automatic irrigation system watered each pot with deionized water once a day. It was programmed for 50ml/day for young plants and achieving 100 ml/day for larger plants. We fertilized the pots on days 15, 45 and 75 with 100 ml Hoagland solution without P and N. On the same days we added 2 ml of full Hoagland solution into the core compartments aiming to stimulate hyphal foraging and growth into the cores.

### Experiment harvest

After 105 days, the experiment was harvested. Pots were harvested one at a time, to avoid delays in freezing the samples for molecular analysis. First, the shoots of the plants were cut and placed in paper bags for later drying and biomass determination. Then, 5 mm of the surface soil of the hyphal compartment cores were removed (to avoid contamination from the surface) and the soil of each core was transferred to a sterile 50ml conical tube and gently homogenized for 30s. From these samples, two aliquots of 2ml soil were transferred to two sterile centrifuge tubes and immediately frozen in liquid N_2_ (to be later used in RNA extraction) and 5ml of soil transferred to a 5ml sterile centrifuge tube and frozen (to be later used for protein extraction). All the spatulas and spoons were treated with 70% ethanol and flamed between samples, and the plastic materials were previously autoclaved. The samples were stored at −80°C. The remaining soil was air-dried for 48h at 40°C and used for the soil aggregation and hyphal length analyses.

### Plant and soil analyses

Plant shoots were dried in an oven for 48h at 60°C and then weighed. We analyzed soil aggregation by measuring the newly formed aggregates and water stable aggregates (WSA), and we also analyzed AMF and non-AMF hyphal length in the soil.

The newly formed aggregates were measured by weighing the dry soil particles >2mm and dividing this value by the total weight of the soil sample. For the WSA measurement we used a wet sieving apparatus (Kemper and Rosenau, 1986). The method was slightly modified as described by Leifheit et al. (2014): 4.0 g of dried soil were rewetted by capillary action and sieved for 3 min using a 250 μm sieve. The coarse matter was separated by crushing the aggregates that remained on the sieve and the small particles washed out through the sieve. Coarse matter and soil were dried at 60°C for 48 h and WSA calculated and corrected for coarse matter.

Hyphal length was determined by extraction from 4.0 g of soil, staining with Trypan blue and followed by microscopy (Jakobsen et al., 1992; Rillig et al. 1999). Dark to light blue stained aseptate hyphae with characteristic unilateral angular projections (“elbows and coils”) were considered mycorrhizal (Mosse, 1959), whereas non-blue stained or blue stained hyphae with regular septation or straight growth were considered non-mycorrhizal (Leifheit et al., 2014). We also counted the number of AMF spores on the slides (Supp. Material 1).

### RNA extraction and sequencing

RNA was extracted from 2g of the frozen soil using the RNeasy PowerSoil Total RNA Kit (Qiagen, Germany). Since an important goal of RNA-seq was to produce a broad and annotated library to allow protein identification, we adopted two library strategies for mRNA enrichment. First, we targeted eukaryotic mRNA enrichment in the samples and used 60% of the volume of each RNA extract for poly- A enrichment using the kit NEXTflex Poly(A) Beads (Perkin Elmer, USA). Secondly, we targeted the transcripts from the full soil diversity using rRNA depletion by hybridization. For that, the four replicates of each treatment were combined in pairs, using 20% of the volume of the RNA extracts from each sample, yielding two samples for each treatment (we pooled samples to increase sequencing depth). The samples were then treated with the kit RiboMinus Bacteria following the manufacturer’s instructions (ThermoFischer, USA).

The 18 mRNA samples, 12 samples poly-A enriched and six samples rRNA depleted, were prepared for sequencing using NextFlex Rapid Directional RNA SeqKit (Perkin Elmer, USA). All the samples were evaluated in an Agilent TapeStation and quantified by QuBit (Invitrogen, USA). The samples of each library were pooled in equimolar ratios for each library strategy. The libraries were then pooled together using a six-fold higher concentration of the rRNA depletion library compared to the poly-A library (since we expected higher diversity for the first). The libraries were sent for sequencing to a commercial company using the Illumina HiSeq × Ten platform. The sequences were deposited at the GenBank server (https://www.ncbi.nlm.nih.gov/genbank - Bioproject number PRJNAxxxxx).

### Soil RNA-seq analysis

The sequences from the HiSeq run were quality checked and demultiplexed using the bcl2fastq software (Illumina, USA). All the sequences were then processed together using the software Trinity and assembled and cleaned of rRNA (Grabherr et al., 2011). The assemblage was annotated by Blast2go (Götz et al., 2008). The sequences were also further classified according to the SEED system from MG-RAST (Wilke et al., 2016). Lastly, the reads of each sample were mapped to the sequences assembled by the software Trinity or to the genome of *R. irregularis* DAO JGI v2 using the software Salmon (Grabherr et al., 2011; Morin et al., 2019). The annotation table and the counting table of the mapped reads were further exported for analysis in R.

### Metaproteomics

Protein extraction was performed based on five grams of frozen soil for all three conditions in biological quintuplicates by the addition of extraction buffer containing 100 mM Tris-HCl pH 8.0, 5% w/v sodium dodecyl sulphate, and 10 mM dithiothreitol. Samples were vortexed and boiled for 10 min at 100°C, centrifuged at 4,000 × g for 10 min and the supernatants were transferred into new protein low-binding tubes. Proteins were then precipitated by the addition of chilled trichloroacetic acid to a final concentration of 20% and kept at −80°C overnight. Proteins were pelleted at 4,000 × g for 30 min, washed twice with cold acetone (−20°C), and air-dried, as described in (Qian and Hettich, 2017).

Protein extracts were reduced and alkylated under denaturing conditions by the addition of 200 μL of a buffer containing 3 M guanidinhydrochlorid, 10 mM tris(2-carboxyethyl)phosphine (TCEP), 40 mM chloroacetamide (CAA), and 100 mM Tris pH 8.5 to prevent proteolytic activities, briefly vortexed and boiled for 10 min at 95°C at 1000 rpm. Pellet disintegration was achieved by a sonicator with five cycles of P150W, C60%, A100% (Hielscher, Teltow, Germany). After centrifugation at 15,000 rcf at 4°C for 10 min, supernatants were transferred into new protein low binding tubes. The lysates were loaded into the preOmics in-stage tip kit cartridges (iST kit 96×, Martinsried, Germany) to remove humics, centrifuged at 3,800 rcf for 3 min, washed with 8 M urea by pipetting up and down ten times and centrifuged again. The lysates were digested and purified according to the preOmics in-stage tip kit (preOmics). In brief, 50 μl lysis buffer was added, diluted with resuspension buffer and digested after 10 min at 37°C at 500 rpm overnight. After several washing steps, peptides were eluted sequentially in three fractions using the SDB-RPS-1 and −2 buffers (Kulak et al., 2014) and the elution buffer provided by preOmics. Eluates were first dried in a SpeedVac, then dissolved in 5% acetonitrile and 2% formic acid in water, briefly vortexed, and sonicated in a water bath for 30 seconds for subsequent analysis on a nanoLC-MS/MS.

### LC-MS instrument settings for shotgun proteome profiling and data analysis

LC−MS/MS was carried out by nanoflow reverse-phase liquid chromatography (Dionex Ultimate 3000, Thermo Scientific, Waltham, MA) coupled online to a Q-Exactive HF Orbitrap mass spectrometer (Thermo Scientific). The LC separation was performed using a PicoFrit analytical column (75 μm ID × 50 cm long, 15 μm Tip ID (New Objectives, Woburn, MA) in-house packed with 3-μm C18 resin (Reprosil-AQ Pur, Dr. Maisch, Ammerbuch-Entringen, Germany). Peptides were eluted using a gradient from 3.8 to 50 % solvent B in solvent A over 121 min at 266 nL per minute flow rate. Solvent A was 0.1 % formic acid and solvent B was 79.9 % acetonitrile, 20 % water, 0.1 % formic acid. An electrospray was generated by applying 3.5 kV. A cycle of one full Fourier transformation scan mass spectrum (300−1750 m/z, resolution of 60,000 at m/z 200, AGC target 1e^6^) was followed by 12 data-dependent MS/MS scans (resolution of 30,000, AGC target 5e^5^) with a normalized collision energy of 25 eV. In order to avoid repeated sequencing of the same peptides a dynamic exclusion window of 30 sec. was used. In addition, only the peptide charge states between two to eight were sequenced.

### Label-Free Proteomics Data Analysis

Raw MS data were processed with MaxQuant software (v1.6.0.1) and searched against the *Rhizophagus irregularis* UP000236242_747089 database, released in May/2019, or the annotated RNA-seq assemblage. A false discovery rate (FDR) of 0.01 for proteins and peptides, a minimum peptide length of 7 amino acids, a precursor mass tolerance to 20 ppm for the first search and 4.5 ppm for the main search were required. A maximum of two missed cleavages was allowed for the tryptic digest. Cysteine carbamidomethylation was set as fixed modification, while N-terminal acetylation and methionine oxidation were set as variable modifications.

The tables generated by MaxQuant were further processed using the online server ANPELA (“analysis and performance assessment of label-free metaproteomes”), data were log transformed and processed by the online pipeline, including missing data imputation by the K-nearest Neighbor algorithm (KNN) and mean normalization (MEA) (Tang et al., 2019).

### Statistical analysis

Analyses were performed in R version 3.5. Plant biomass, water stable aggregates and hyphal length were analyzed by ANOVA, followed by the Tukey posthoc test. We also correlated variables using Spearman’s rank correlation coefficient (*p*). The distribution of the data and effect sizes were analyzed by the R-DBR package (SI Figure S1).

The tables previously obtained from the RNA-seq and proteomics were aggregated with the SEED functions for a first explanatory analysis. Differences between the samples were analyzed by cluster analysis. The taxonomy obtained by the Blast2go annotation was used in the diversity analysis and RDA using R-Vegan. The data were further analyzed by R packages DeSeq2/Alx2 and the online server GMine (http://cgenome.net/gmine) (Quinn et al., 2018). Since both RNA-seq data sets were analyzed according to the pool of identified sequences, we treated them as compositional data, considering the centered log-ratio of the values of each gene/protein for each sample (Gloor et al., 2017). We report the log-ratio fold change of the gene/proteins and Welch’s t-test with Benjamini-Hochberg corrected statistical probabilities, calculated by the R-packages DeSeq2 and Alx2 or the online server Gmine.

## 3. Results

We evaluated the effects of *R. irregularis* inoculation compared to control soil. Plants were harvested at 105 days, but no difference was observed in the shoot plant biomass (Figure 1 A-B and SI S1).

**Figure 1.**
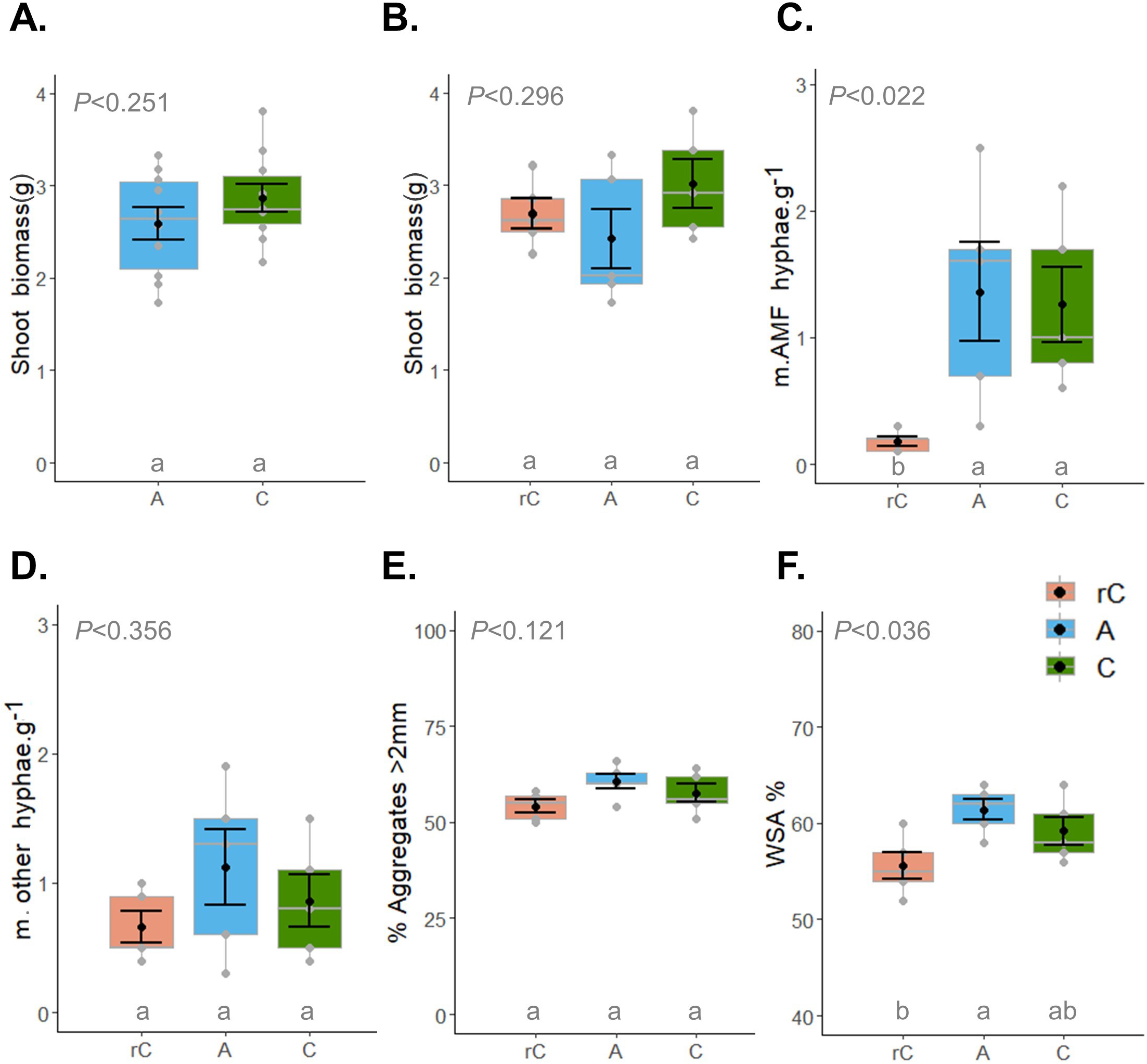
A. Dry shoot biomass of clover inoculated with the AMF *R. irregularis* (A) and the non-inoculated controls (C) (n=10). B. Shoot biomass for the main treatments A, C and including the rotated core control (rC) (n=5). The full comparisons with all replicates are available in Supp. Material 1. C. AMF hyphal length (in m g^−1^ of soil). D. Hyphal length of other fungi (in m g^−1^ of soil). E. New aggregates > 2mm (%) F. Water stable aggregates (%). The ANOVA *P*-value results are reported on the bottom of each graph and small letters below the treatment codes, when different, indicate statistical differences by the Tukey posthoc test (*P*<0.05). The gray dots on the box-plots indicate the individual value of each replicate. The dark dot indicates the mean and error bars the standard error.

We further evaluated the soil compartments that were divided in three treatments with five replicates each: 1) control with static cores (C), 2) control with rotated cores (rC) and 3) cores of soil inoculated with the AMF *R. irregularis* (A). First, we detected that the C and A cores were colonized by AMF, while very few AMF hyphae were observed in the rC (Figure 1.C). We also detected colonization by non-AMF hyphae in all the cores, but there was no difference (*P*=0.356) among treatments (Figure 1 D). Secondly, we detected increases in soil aggregation. At the beginning of the experiment, all soil was sieved to 2 mm and we observed at the harvest time 57 ± 4 % new formed aggregates, however no strong differences were observed across treatments (*P*=0.121). We also observed increases in WSA during the experiment, from 46 % ± 3 at the beginning, increased to 56-61 % after 105 days (Figure 1.E.). At 105 days we found higher soil aggregation in the cores inoculated with *R. irregularis* in relation to the rotated cores (*P*=0.036) (Figure 1.D-E). We also detected a correlation between AMF hyphal length and WSA across the treatments (*p=*0.67, *P*<0.01), as also between non-AMF hyphae and WSA (*p=*0.60, *P*<0.01).

### Effects of AMF on soil community metabolism

The samples were further analyzed in terms of metatranscriptomics (Figures 2-4). The HiSeq sequencing run yielded 180 million reads (SI Figure S2). From that, for the rRNA depletion libraries we recovered an average of 6 million reads for the C treatment, 2.5 million for rC and 6.5 million for A. For the poly-A libraries we recovered an average of 21.5 million reads for the C treatment, 30 million for rC and 25 million for A. For the Poly-A enrichment library, around of 50% of the transcripts were still from bacteria (Figure 4). This indicates that Poly-A selection from soil is challenging, since most of the transcripts in soils are generally from bacteria (Mendes et al., 2018a; Schlüter et al., 2019). The method could be improved in future by, for example, by repeating the poly-A selection for several times. Based on the Blast2go annotation, from the poly-A library, we were able to recover around 0.47% of reads of AMF (*Glomeromycota*) for C, 0.36% for rC and 0.46% for A (SI Figure S2). For the rRNA depletion library we were able to recover 0.45% AMF reads for C, 0.32% reads for rC and 0.42% for A.

**Figure 2.**
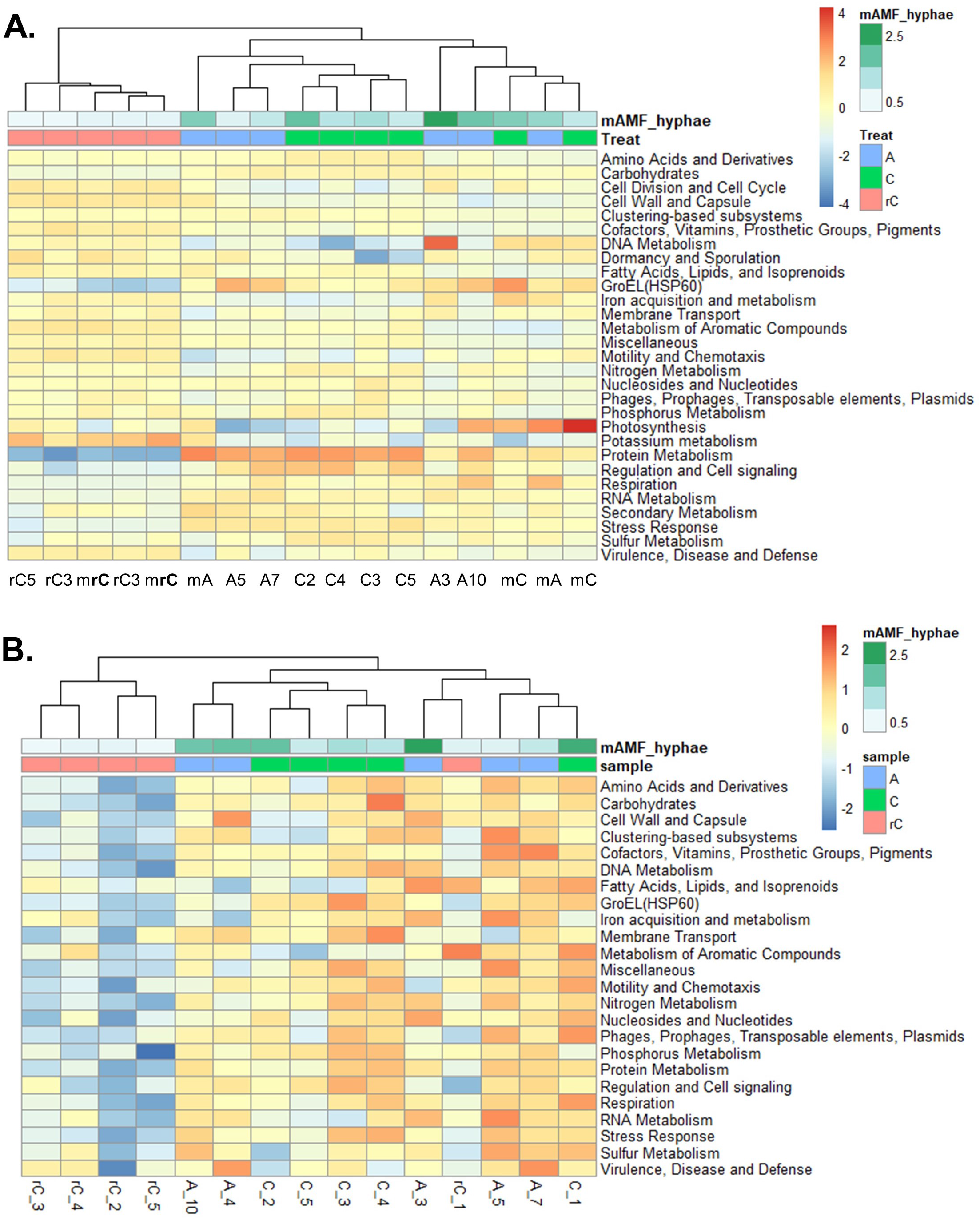
Metabolic functions in soil clustered by the MG-RAST SEED classification accessed by soil metatranscriptomics **(A)** and metaproteomics **(B)**. The sample codes are: static soil core inoculated with the AMF *R. irregularis* (A), non-inoculated static soil core (C), and non-inoculated rotated soil core (rC). For the RNA samples, the prefix **m** indicates the mRNA libraries enriched by rRNA depletion, the other samples without prefix indicate the poly-A selection libraries.

The transcript analyses indicated several differences in gene expression related to C, N, P and respiration processes clustered by the MG-RAST SEED terms (Figure 2-3). From the 616,863 genes assembled by Trinity, 161,702 could be annotated to the 29 SEED categories “level 1”. We further segregated the “GroEL/HSP60” category (SEED level 2) from the category “protein” in SEED level 1, to better test our third hypothesis about HSP60. The first exploratory analysis, the heat map and cluster analysis, indicated that the samples were distinctly clustered between the rotated core and the other treatments with high abundance of AMF hyphae (Figure 2.A). Some clustering was observed differentiating the static core control and the *R. irregularis* treatment (Figure 2.A). Notice that the heat map scale is not about negative or positive gene expression, but it is based on the scaled centered log-ratio (CLR) transformation recommended for compositional data and reflects the relative abundance of each metabolic category in each sample. From the 29 functional categories, 15 categories were associated by a network analysis to the presence of AMF hyphae in soil (Figure 3). The analyses indicated that *R. irregularis* inoculation had increased expression of genes related to RNA metabolism, respiration and GroEL/HSP60. The control sample, that also had AMF hyphae, had intermediate values between the inoculated and the rotated core treatments and was related with C, N, P and S metabolism. The rotated control was related mostly with cell division, mobility, fatty acids, and secondary metabolism (Figure 3). We found a correlation between AMF hyphal length and transcripts related to respiration (r=0.66, *P*=0.028), but no strong evidence for a correlation between hyphal length and total GroEL/HSP60 transcripts (r=0.47, *P*=0.14).

**Figure 3.**
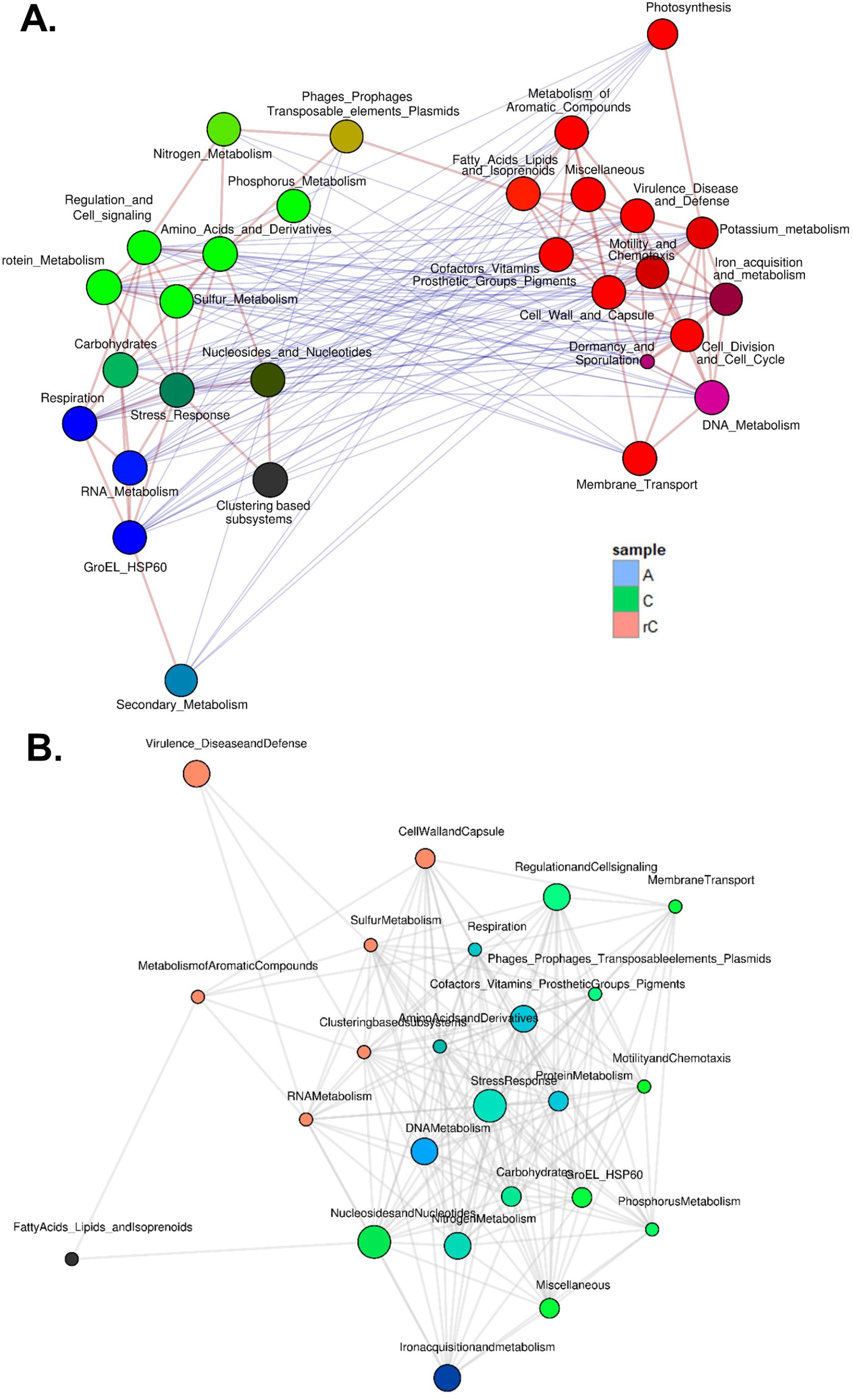
The network connectivity of soil metabolic functions (SEED level 1) across treatments for the metatranscriptome (**C.**) and the metaproteome (**D**.). In blue the treatment inoculated with the AMF *R. irregularis* (A), in green the control soil (C) and in salmon the control with rotated soil cores (rC).

**Figure 4.**
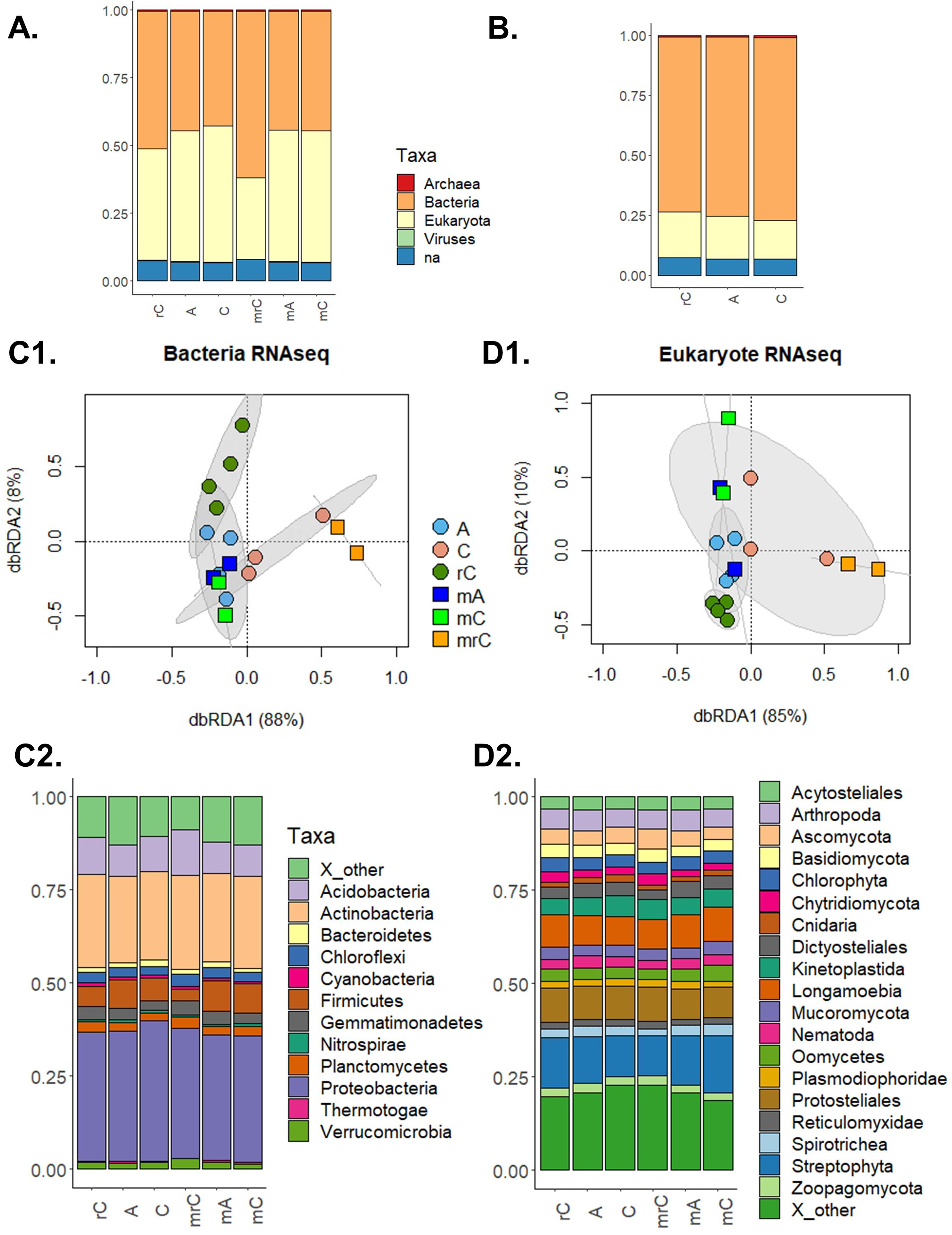
Changes in the soil microbial community structures based on the taxonomic annotation of the sequences. On top are the general high order taxonomy for the RNA-seq data (**A.**) and for the proteome data (**B**.) (for details on the proteome see Supp. Figure 4). In the middle are the microbial community structures accessed by the redundancy analysis (RDA) based on the RNA-seq for *Bacteria* (**C1.**) and for *Eukaryota* (**D1.**). Below are the phyla relative abundance of *Bacteria* (**C2**) and *Eukaryota* (**D2**) across the treatments. The sample codes are: static soil core inoculated with the AMF *R. irregularis* (A), non-inoculated static soil core (C), and non-inoculated rotated soil core (rC). For the RNA samples, the prefix **m** indicate the mixed libraries made using the rRNA depletion.

For the proteomic data, we identified 1336 proteins using the RNA-seq as reference, and the data normalized by ANPELA indicated a strong separation of the samples by treatment (SI Figure S3), and a volcano plot identified 103 proteins that were more abundant in C and A than in the rotated cores (Supp. Figure 3). From all the proteins, 398 were identified at MG-RAST SEED categories (Figure 2.B), and, again, there was a differentiation between rotate cores to the control and soil inoculated with *R. irregularis*. Considering these 398 proteins, there was a decrease in 19 SEED categories in the rotated cores (Figure 3.B) and thus the network analyses were strongly influenced by the non-rotated treatments, that can be observed by the color of the network nodes, indicating mainly the non-rotated treatments (Figure 3.D). GroEL/HSP60 was more abundant in the soil cores inoculated with *R. irregularis*. Using the *R. irregularis* genome as reference, we identified 51 proteins and most of them were proteins related to protein metabolism, including heat shock proteins (we report on this separately in the section about HSP).

### Effects of AMF on soil community structure

The metatranscriptomic data were then analyzed taxonomically. From the 616,863 genes assembled by Trinity, 199,566 were annotated by Blast2go. All the samples were dominated by Bacteria (Figure 4). The rRNA depletion library had approx. 45% of Bacteria for the A and C treatments and approx. 65% in the rC (Figure 4.A). For the poly-A libraries, it was also approx. 45% of Bacteria for the A and C treatments, but approx. 50% of *Bacteria* in the rC. The microbial community structure was further investigated by RDA (Figures 4. C1 and D1). For both, *Bacteria* and *Eukaryota*, the samples were better clustered by the libraries made from the mRNA enrichment by rRNA depletion libraries (“m” code in the figures). For the depletion libraries, the rotated cores had a distinct community structure when compared to the static-control soil and the soil inoculated with *R. irregularis* (perMANOVA, *P*<0.001 for both, *Bacteria* and *Eukaryota*). However, clustering of the samples from the poly-A libraries was not strongly supported (perMANOVA, *P*<0.12 for *Bacteria* and P<0.20 for *Eukaryota*).

The samples were further analyzed in detail in terms of the composition of *Bacteria* and *Eukaryota* sequences. The RNA-seq data indicated that all the samples were dominated by *Actinobacteria* and *Proteobacteria* (no difference among treatments was observed; *P*=0.2), but a reduction of *Firmicutes* (Aldex2, *P*=0.052) was observed in the rotated cores (Figure 4.C2). Considering *Eukaryota*, the samples were dominated by *Longamoebia*, *Protosteliales* and *Streptophyta* (Figure D2). *Fungi* accounted for around 15% of the *Eukaryota* reads, and *Glomeromycota* for around 0.4% of the reads (SI Figure S2).

### Effects of AMF on transcripts/proteins potentially linked to soil aggregation

We screened for genes/proteins potentially related to soil aggregation. We explored a list of genes/proteins including genes related to EPS metabolism (Cania et al., 2019), such as polysaccharide export outer membrane (Wza), capsular polysaccharide export system (Kps) and lipopolysaccharide export proteins (Lpt). We did not find differences for most of the genes/proteins, but found in the metatranscriptome that Lpt (Figure 5A) was more abundant in rotated core samples (z-score 0.6-0.9), when compared to the control and *R. irregularis* inoculated treatments (*P*<0.001).

**Figure 5.**
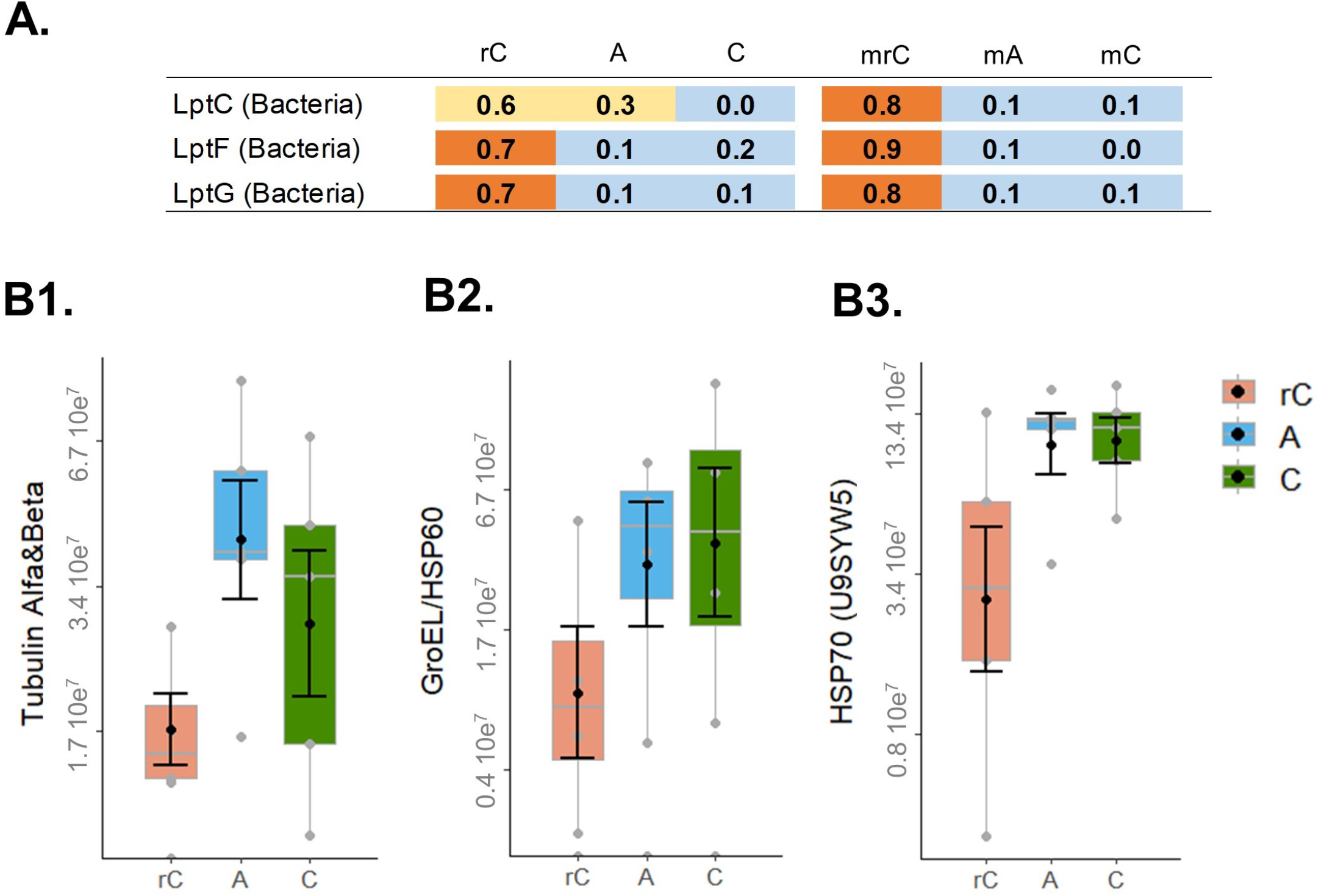
The genes and proteins potentially related with soil aggregation. The panel on top show the z-score of Lpt genes across the samples (**A.**). Below is the quantification of proteins identified in relation to the *R. irregularis* proteome database. The MS intensity signal for Tubulin (Alfa+Beta) (**B1.**), GroeEL/HSP60 (**B2.**) and HSP70 (uniprot U9SYW5) (**B3.**). The sample codes are: static soil core inoculated with the AMF *R. irregularis* (A), non-inoculated static soil core (C), and non-inoculated rotated soil core (rC).

We further evaluated HSP60/GroEL, which was very abundant across all treatments, especially in the cores with higher abundance of AMF hyphae (Figure 2). In all treatments, at the RNA level, HSP60/GroEL was predominantly of bacterial origin, particularly from *Acidobacteria* and *Proteobacteria* (SI Figure S5 A1-2). Of the few reads of eukaryotic origin, the great majority was from *Basidiomycota*, with the exception of the rotated cores, which had a high abundance of reads from *Longamoebia* (SI Figure S5 A3). In the proteome, the vast majority of identified HSP60/GroEL proteins came from *Bacteria* (app. 99% across all treatments), and they were predominantly from *Acidobacteria* and *Proteobacteria* (SI Figure S5 B).

We found a correlation between the total HSP60/GroEL with the AMF hyphal length for the total metaproteome (r=0.57, *P*=0.026), but little evidence for the total metatranscriptome (r=0.47, *P*=0.14). However, as described before, most of these HSP60/GroEL came from *Bacteria* (SI Figure S5 A1). We observed no correlation between total HSP60/GroEL and WSA (all had *P*>0.2).

We further looked at the proteins identified from *R. irregularis* in the metaproteome and there were no differences in HSP60/GroEL (*P*=0.21) across the treatments (Figure 6). We compared this result with tubulins (Alfa+Beta) (*P*=0.08), which are essential proteins related to fungal growth, and observed a very similar MS signal intensity to the HSP60/GroEL protein. We screened the other HSPs from *R. irregularis* and detected also little evidence for differences across treatments for two other HSP, the HSP70 (Uniprot A0A2H5SZG6, *P*=0.14) and HSP70 (Uniprot U9SYW5, *P*=0.07). None of these proteins were correlated with AMF hyphal length or WSA (all had *P*>0.2). These results also indicated that *R. irregularis* transcripts and proteins were a small proportion of the full microbial community (we estimated AMF being between 0.3-0.5% of the active community, SI Figure S1).

## 4. Discussion

In this study, we investigated the effects of AMF on soil microbial communities using soil RNA-seq and metaproteomics. We detected that AMF drove the active microbial community as well as transcripts and proteins related to metabolic functions. First, we showed that the presence of AMF induced changes in transcripts and proteins in soil related to several metabolic processes, including protein metabolism (N cycle) and respiration (C cycle). Secondly, we showed that *R. irregularis* induces changes in the soil microbial community, which was largely dominated by *Bacteria*. And thirdly, we detected transcripts and proteins that could be related to soil aggregation.

Previous knowledge about AMF affecting soil functions was obtained mostly by inoculation studies, or by quantifying AMF hyphae in soils and correlating them with specific functions, such as nitrogen uptake or soil aggregation (Powell and Rillig, 2018; Hestrin et al., 2019; Rillig et al., 2019). To the best of our knowledge, this was the first attempt to experimentally assess metatranscriptomes and metaproteomes to estimate the contribution of AMF to the metabolism of the entire soil community. We find that only 0.3-0.5% of the transcripts/proteins were of AMF origin and our data indicate that AMF induced significant shifts in soil community metabolism when compared to the rotated core control (Figures 2 and 3).

Our results bridge between microbial ecology and ecosystems functioning. The data suggest that the AMF hyphosphere drive an active community related to soil respiration and nutrient cycling, potentially improving nutrient mineralization from soil organic matter and nutrient supply to plants. Previous studies also presented several lines of evidence that AMF influence the microbial community and that these fungi are important players in the carbon cycle, P and N dynamics and soil respiration (Nottingham et al., 2010; Zhang et al., 2016; Powell and Rillig, 2018; Hestrin et al., 2019). Interestingly, previous studies have suggested that AMF are associated with specific bacterial communities and that these communities have complementary functional roles to the AMF (Battini et al., 2016; Turrini et al., 2018). Our data corroborate these findings in a general way, since the community had increased expression of genes/proteins related to general soil processes in the presence of the AMF hyphae (Figures 2 and 3).

Thus, we present evidence that arbuscular mycorrhizal fungi (AMF) select for beneficial microbial communities in their hyphosphere. We suggest two possible mechanisms to be tested in future: 1) the AMF hyphae produce exudates that select a beneficial microbial community, analogous to reports for plant roots (Baltrus, 2017); or, 2) AMF hyphae compete with the microbial community for available nutrients and consequently induce the community to mineralize nutrients from soil organic matter (Hestrin et al., 2019).

Moreover, our study showed that the soil communities were dominated by transcripts and proteins of bacterial origin (Figure 4). We observed that the samples were dominated by *Proteobacteria* and had an increased relative abundance of *Firmicutes* in the hyphal compartments, and both groups were previously reported to be predominantly associated with AMF (Battini et al., 2016; Turrini et al., 2018). Interestingly in the rotated cores there was an increased expression of bacterial *lpt* genes (lipopolysaccharide export proteins). These genes can be related to soil aggregation and we speculate that these could be related to soil aggregation in the rotated cores, since soil aggregates increased from the beginning of the experiment to the harvest (Cania et al., 2019).

Lastly, our data showed a high abundance and diversity of HSP60 (GroEL) across the soil samples detected by RNA-seq and by proteomics. This can be explained in two ways; first, this chaperon occurs across all living organisms and it has essential functions in protein folding and cell metabolism, in addition to moonlight functions (Henderson et al., 2013). Second, for the proteome samples, our method included the boiling of samples and this can likely select for thermostable proteins, such as HSPs (Rosier et al., 2006; Gillespie et al., 2011). Furthermore, HSP60 abundance may not only reflect cell abundance, but also the metabolic status of cells (Henderson et al., 2013). Moreover, our data clearly showed the high complexity of a soil community sample and that HSP60 is highly abundant and diverse in soil. We also observed that at nucleotide level the HSP from *R. irregularis* are divergent enough to be differentiated from the community sequences (Magurno et al., 2019), but due codon redundancy they are highly conserved at the amino acid level. This implies that during processing for proteomics the HSP60 could potentially be cleaved in positions to overlap MS peptides spectra of HSP60/GroEL from other members of the microbial community, and theoretically causing noise in the quantification of HSP60 from AMF. Previous studies using the monoclonal antibody MAb32B11 for *R. irregularis* already showed that HSP60 quantifications in soil are also difficult due to the complex composition of organic material in soil and could make quantifications imprecise (Rosier et al., 2006). Our new findings also bring additional evidence to avoid quantifying GRSPs as a proxy to quantify AMF proteins in soil (Rosier et al., 2006; Gillespie et al., 2011). Nevertheless, we are optimistic that future works using high-resolution proteomics, perhaps in simpler soil communities, will be able to identify specific proteins from AMF in soil and potentially correlating them with ecosystems functions, such as soil aggregation.

## 5. Conclusions

We provide evidence that the AMF hyphosphere drives an active community related to soil respiration and nutrient cycling, potentially improving nutrient mineralization from soil organic matter and nutrient supply to plants. Overall, our results show that AMF have an intense effect on the soil microbial community and on the metabolic pathways related to soil functions and nutrient dynamics. These findings contribute to a better understanding of the role of AMF in soil and how they drive soil metabolism and microbial community structure.

## 6. Acknowledgments

We acknowledge Alexander-von-Humboldt Foundation and CAPES Foundation for a postdoctoral grant awarded to DRL, and Freie Universität Berlin and the Max Planck Society for funding this work.

## Supporting Information

**Figure S1** Cumming plots for the classical variables evaluated in the experiment. **A.** Clover dry shoot biomass. **B.** AMF hyphae length (in m.g^−1^ of soil). **C.** Hyphae length of other fungi (in m.g^−1^ of soil). **D.** AMF spores observed during the hyphae length measurements **E.** New aggregates > 2mm (%) **F.** Water stable aggregates (%). The codes are: A, soil inoculated with the AMF *R. irregularis*, C, control soil (static core), and rC, the rotated core soil.

**Figure S2** Summary of the RNA-seq results. In panel **A.** the read counts across the treatments; in **B.** Effect size and Volcano plot generated by Aldex2 showing all the transcripts as dots and in light colors are the transcripts that had statistical significant changes in relation to the rotated core (*P*<0.05).

**Figure S3** Proteome data after the ANPELA server processing. A. Heatmap of all the identified proteins according to treatments (ANPELA normalized values). The sample codes are: static soil core inoculated with the AMF *R. irregularis* (A), non-inoculated static soil core (C), and non-inoculated rotated soil core (rC). B. A volcano plot showing significantly regulated proteins between the AMF treatments and the rotated core control (*P*<0.01).

**Figure S4** The relative abundance of the all proteins identified by the taxonomic annotation for *Bacteria* (**A.**) and *Eukaryota* (**B.**) across the treatments. The sample codes are: static soil core inoculated with the AMF *R. irregularis* (A), non-inoculated static soil core (C), and non-inoculated rotated soil core (rC).

**Figure S5** The relative abundance of the GroeL/HSP60 reads of the RNAseq based on the high order taxonomy (**A1.**) and according to *Bacteria* (**A2.**) and *Eukaryote* (**A3.**) phyla. On the bottom right is the relative abundance of the GroeL/HSP60 proteins across the proteomes (**B**). The sample codes are: static soil core inoculated with the AMF *R. irregularis* (A), non-inoculated static soil core (C), and non-inoculated rotated soil core (rC). For the RNA samples, the prefix m indicate the mixed libraries made using the rRNA depletion.

