## Supporting Information for "The effects of arbuscular mycorrhizal fungi (AMF) and *Rhizophagus irregularis* on soil microorganisms assessed by metatranscriptomics and metaproteomics"

Figure. S1
Figure. S2
Figure. S3
Figure. S4
Figure. S5

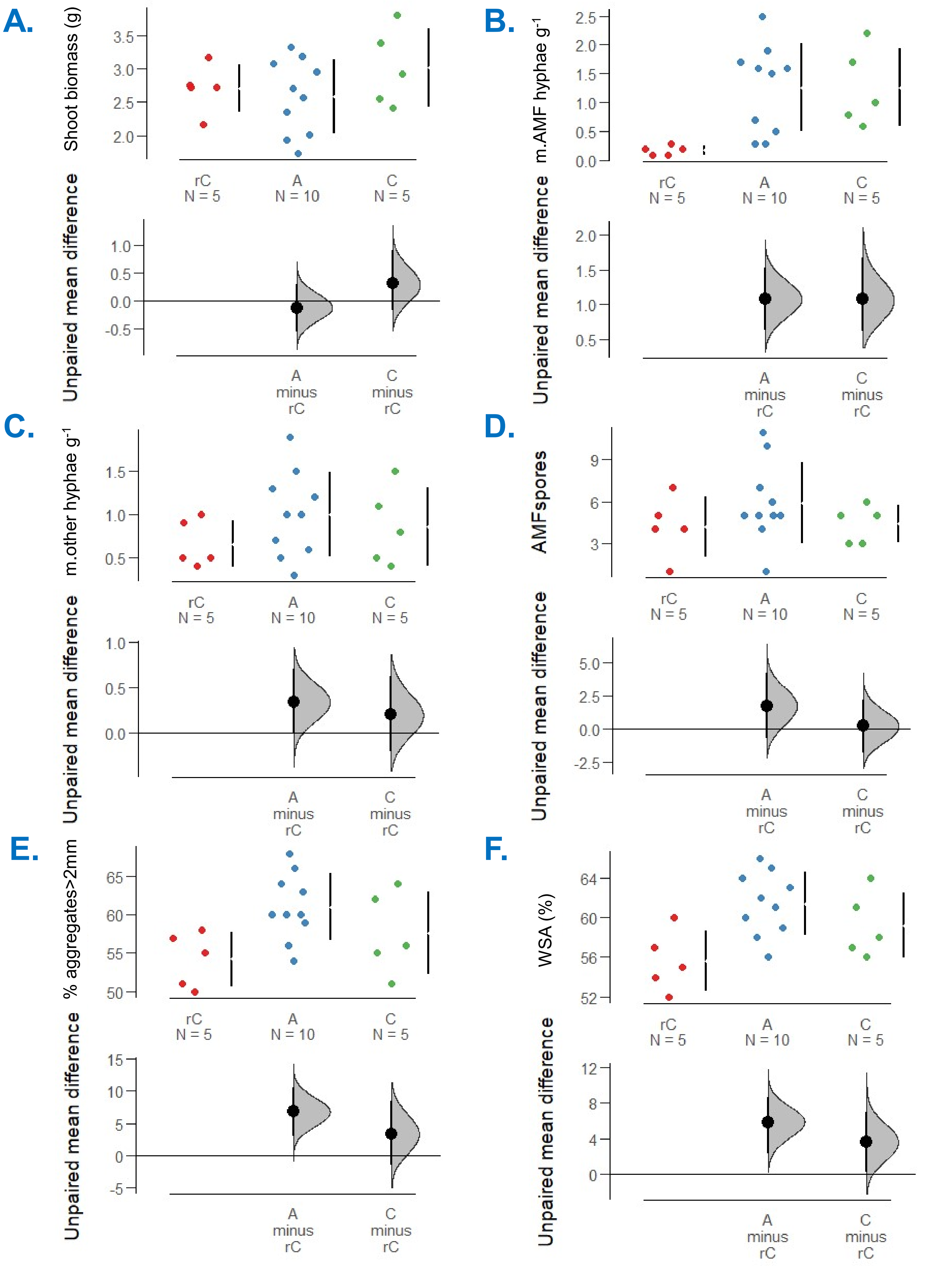


**Figure S1** Cumming plots for the classical variables evaluated in the experiment. **A.** Clover dry shoot biomass. **B.** AMF hyphae length (in $m.g^{-1}$ of soil). **C.** Hyphae length of other fungi (in $m.g^{-1}$ of soil). **D.** AMF spores observed during the hyphae length measurements **E.** New aggregates > 2mm (%) **F.** Water stable aggregates (%). The codes are: A, soil inoculated with the AMF *R. irregularis*, C, control soil (static core), and rC, the rotated core soil.

## A.

| Sample | reads_HiSeq | reads_rRNA | Count_Table | Counts_match_BlastX | Glomeromycota (BlastX) | Reads maped in R.i. genome |
|---|---|---|---|---|---|---|
| C | 4,914,820 | 3,877,758 | 1,037,062 | 337,220 | 0.48% | 162,031 |
| C | 5,347,198 | 4,307,960 | 1,039,238 | 343,148 | 0.45% | 182,620 |
| C | 14,024,377 | 10,965,872 | 3,058,505 | 1,039,768 | 0.48% | 452,040 |
| C | 1,571,230 | 117,913 | 1,453,317 | 536,493 | 0.45% | 252,811 |
| rC | lost library | - | - | - | - | - |
| rC | 2,807,020 | 93,360 | 2,713,660 | 1,107,787 | 0.40% | 703,010 |
| rC | 3,316,560 | 218,954 | 3,097,606 | 1,182,783 | 0.30% | 234,867 |
| rC | 2,576,780 | 530,425 | 2,046,355 | 858,720 | 0.40% | 459,661 |
| A | 21,254,765 | 19,017,341 | 2,237,424 | 967,890 | 0.39% | 818,603 |
| A | 2,999,000 | 68,384 | 2,930,616 | 1,162,604 | 0.41% | 551,553 |
| A | 3,049,410 | 211,982 | 2,837,428 | 1,076,956 | 0.48% | 638,961 |
| A | 1,862,270 | 367,067 | 1,495,203 | 602,656 | 0.56% | 463,477 |
| mA | 30,931,703 | 25,056,043 | 5,875,660 | 2,121,957 | 0.42% | 1,129,739 |
| mA | 20,486,300 | 17,869,783 | 2,616,517 | 1,103,289 | 0.42% | 856,599 |
| mC | 21,957,137 | 19,765,963 | 2,191,174 | 952,708 | 0.50% | 862,337 |
| mC | 22,531,373 | 20,495,575 | 2,035,798 | 860,441 | 0.40% | 943,152 |
| m**rC** | 42,895,361 | 32,195,165 | 10,700,196 | 4,041,269 | 0.33% | 659,005 |
| m**rC** | 19,147,946 | 13,850,654 | 5,297,292 | 1,969,422 | 0.32% | 148,856 |

## B.

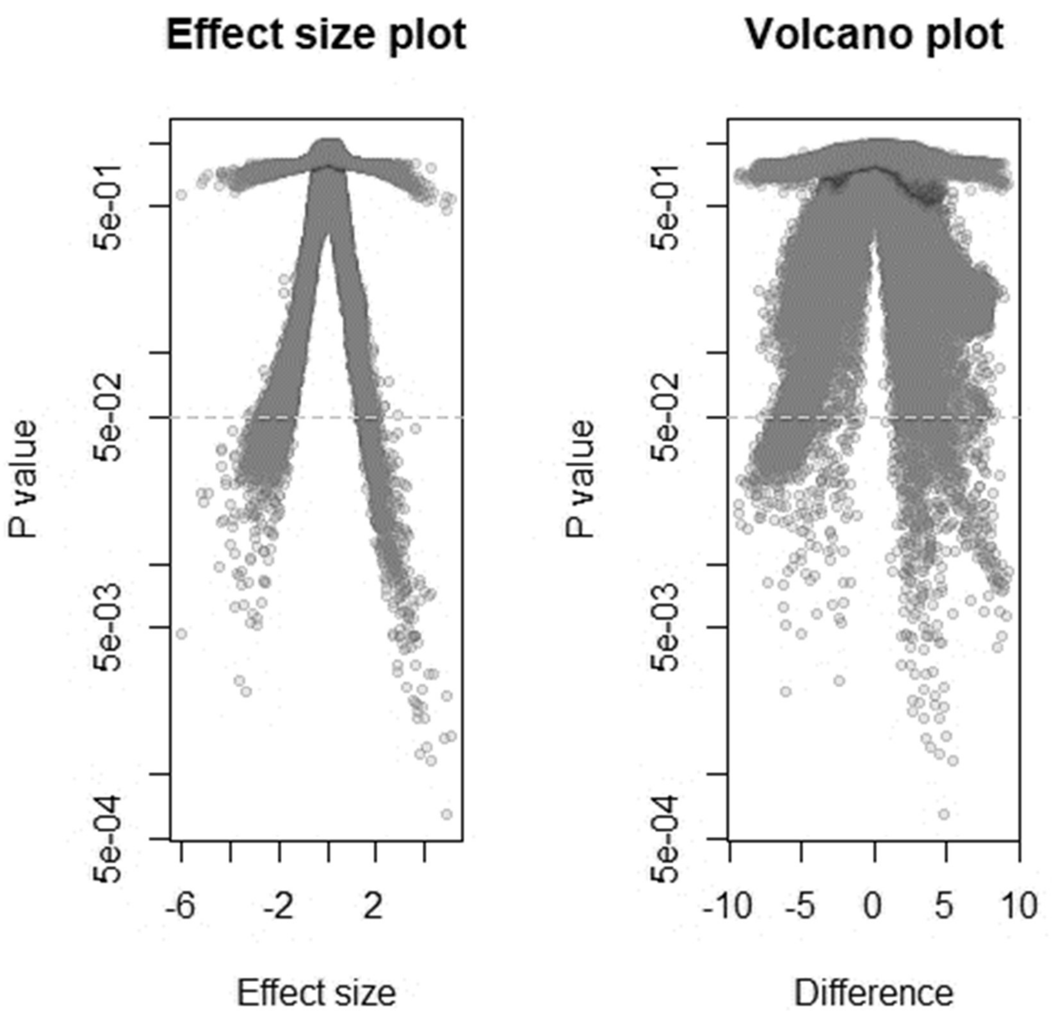


**Figure S2** Summary of the RNAseq results. In panel **A.** the read counts across the treatments; in **B.** Effect size and Volcano plot generated by Aldex2 showing all the transcripts as dots and in light colors are the transcripts that had statistical significant changes in relation to the rotated core (*P*<0.05).

**A.**

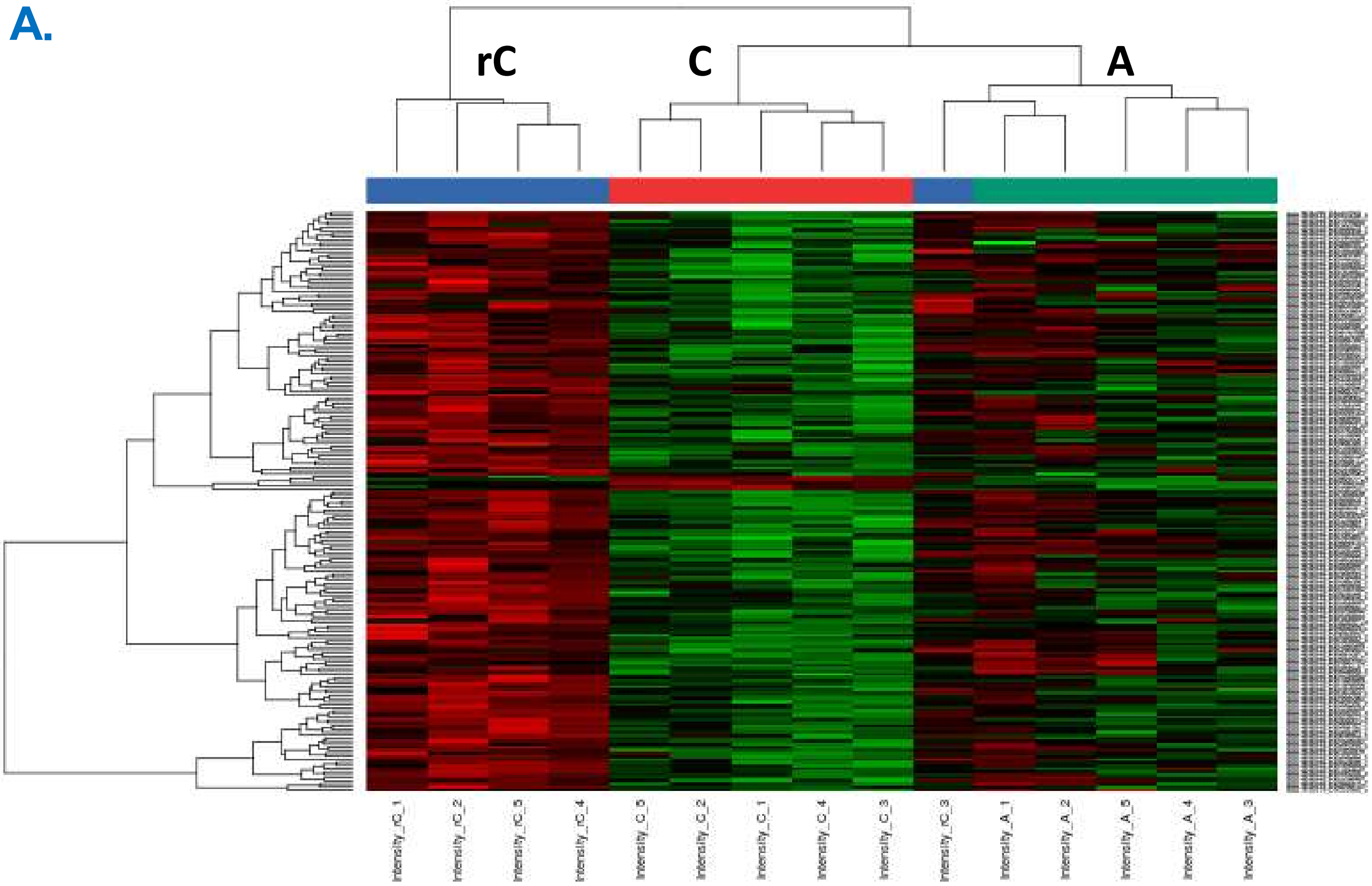


**B.**

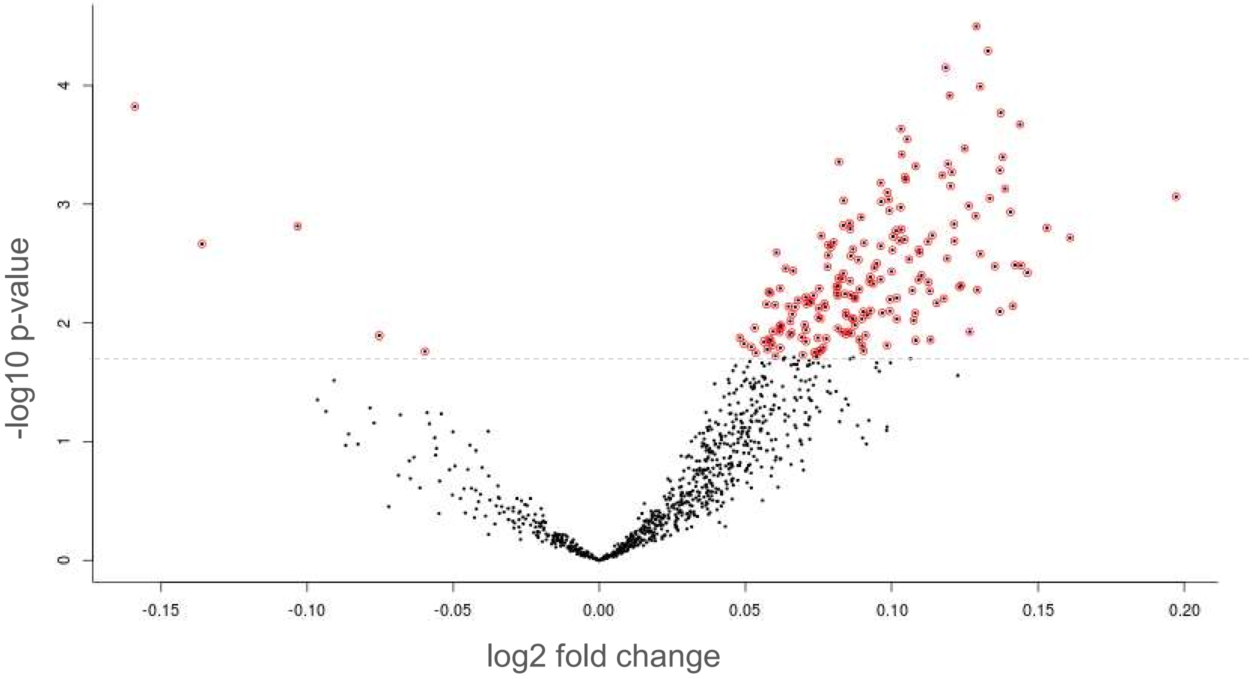


**Figure S3** Proteome data after the ANPELA server processing. **A.** Heatmap of all the identified proteins according to treatments (ANPELA normalized values). The sample codes are: static soil core inoculated with the AMF *R. irregularis* (A), non-inoculated static soil core (C), and non-inoculated rotated soil core (rC). **B.** A volcano plot showing significantly regulated proteins between the AMF treatments and the rotated core control (*P*<0.01).

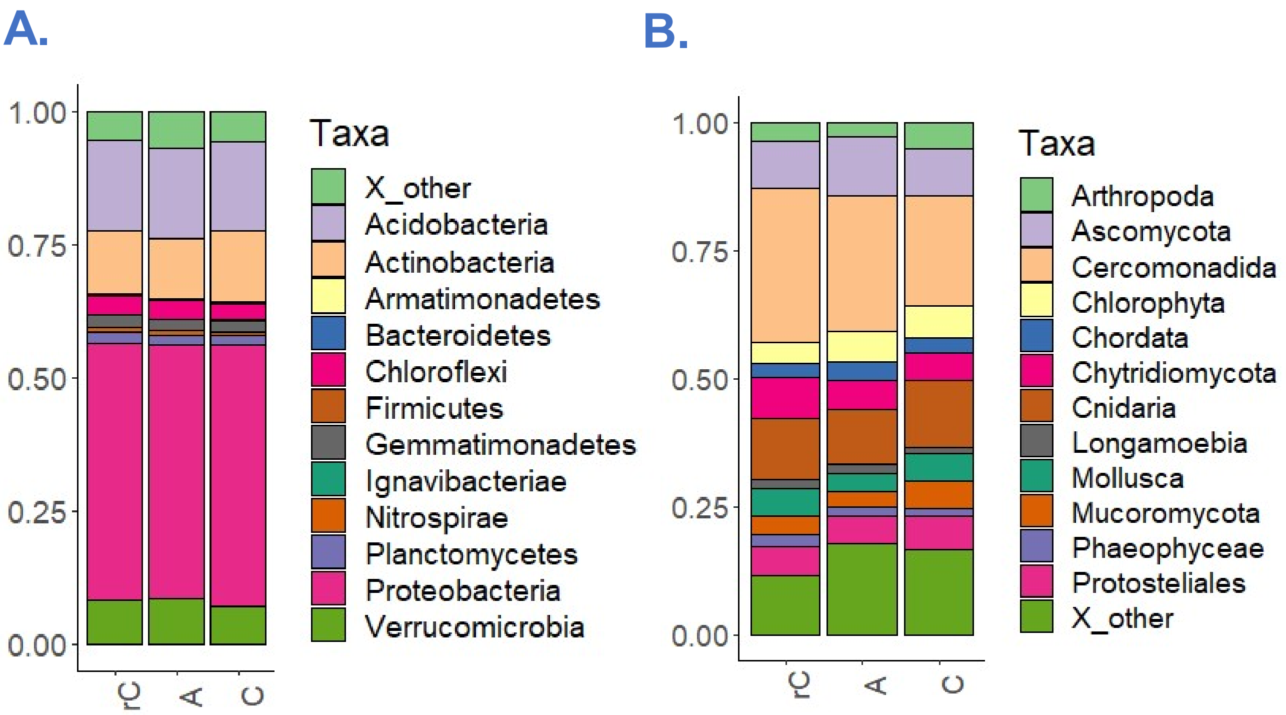


**Figure S4** The relative abundance of the all proteins identified by the taxonomic annotation for *Bacteria* (**A.**) and *Eukaryote* (**B.**) across the treatments. The sample codes are: static soil core inoculated with the AMF *R. irregularis* (A), non-inoculated static soil core (C), and non-inoculated rotated soil core (rC).

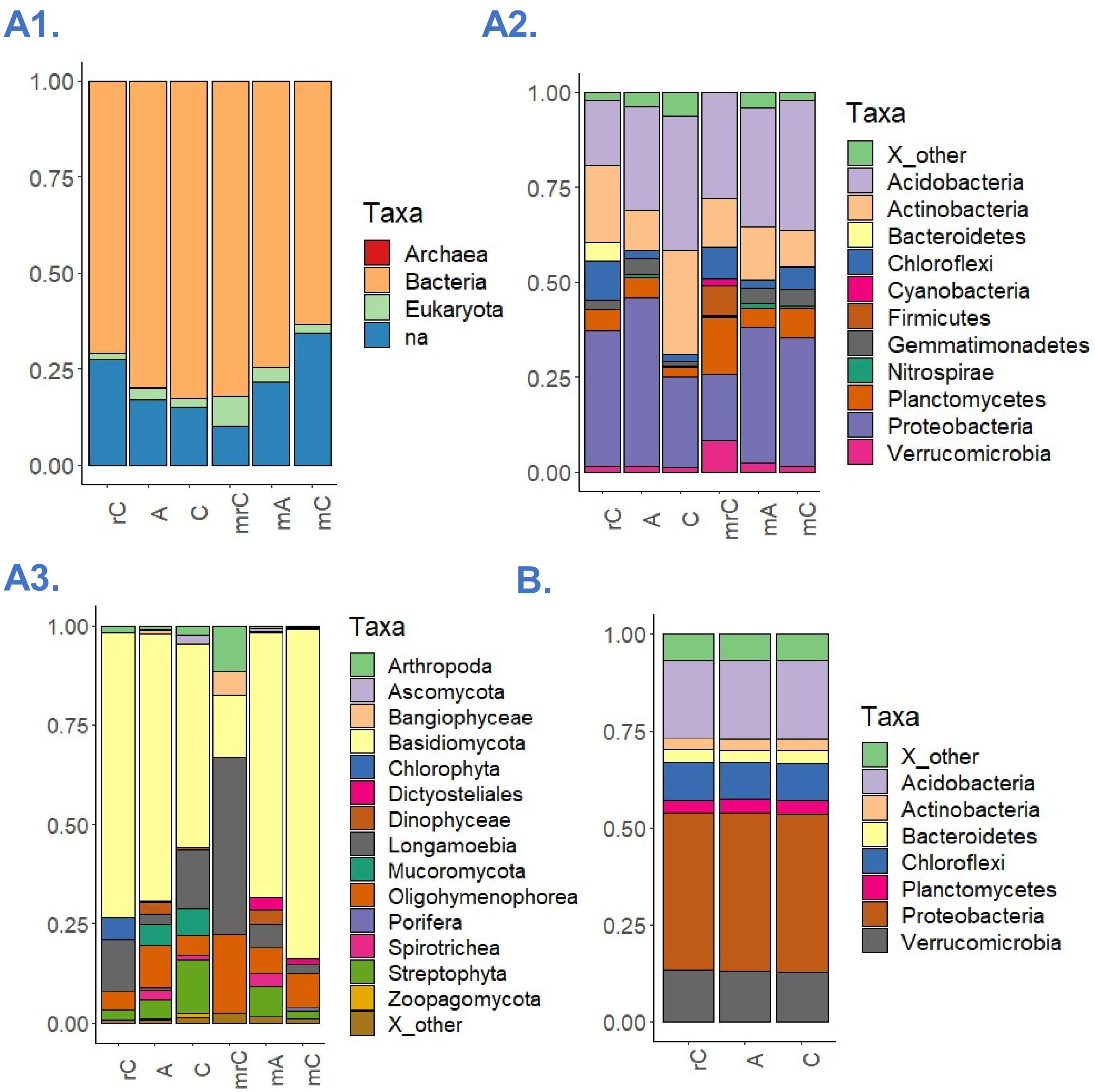


**Figure S5** The relative abundance of the GroeL/HSP60 reads of the RNAseq based on the high order taxonomy (**A1.**) and according to *Bacteria* (**A2.**) and *Eukaryote* (**A3.**) phyla. On the bottom right is the relative abundance of the GroeL/HSP60 proteins across the proteomes (**B**). The sample codes are: static soil core inoculated with the AMF *R. irregularis* (A), non-inoculated static soil core (C), and non-inoculated rotated soil core (rC). For the RNA samples, the prefix m indicate the mixed libraries made using the rRNA depletion.
